# Impact of Donor Individuality, Temporal Variation, and Culture Medium Type on Microbiota Composition and Metabolic Activity in Human Fecal Batch Culture

**DOI:** 10.1101/2023.08.07.552316

**Authors:** Zhuang Liu, Jacoline Gerritsen, Hauke Smidt, Erwin G. Zoetendal

## Abstract

Fecal batch culture (FBC) studies often rely on a single fecal sample collection and the use of one type of medium for cultivation, bringing challenges to the interpretation of results and the comparison between studies. This study investigated the impact of donor individuality, temporal variation and culture medium type on microbiota composition and metabolic activity in an FBC setting with the fiber polydextrose (PDX) as carbon and energy source. FBCs were inoculated with fecal microbiota from three healthy donors sampled at three different days (day 1, 2 and 30), using either basal or rich culture medium with PDX as carbon source. Microbiota composition and metabolic activity were determined after 0, 6, 12, and 24 h of incubation. Microbiota composition variation explained by donor individuality dropped from 51% to 16% during incubation, while that explained by medium and PDX supplementation increased from 0% to 17% and 20%, respectively. Independent of the medium, the genera *Erysipelotrichaceae* UCG-003, *Blautia* and *Fusicatenibacter* were stimulated by PDX supplementation. In basal medium *Bacteroides* and *Anaerostipes* grew better, whereas *Bifidobacterium*, *Faecalibacterium* and *Megasphaera* grew better in rich medium. Metabolite variation was explained up to 50% by PDX supplementation during incubations, with butyrate being produced at the highest concentrations among all metabolites. Temporal variation explained less than 3% of the variation in both microbiota and metabolite composition. In conclusion, in this study donor individuality had the most profound impact on microbiota succession while medium and PDX supplementation had larger impacts on metabolic activity in FBCs.

**IMPORTANCE:** FBCs or other *in vitro* models are often chosen to assist in obtaining mechanistic insights complementing *in vivo* microbiome observations by mimicking the colonic fermentation. However, FBCs are prone to a variety of factors such as the individuality of feces donor, temporal variation in microbiota composition within the individual, and cultivation medium. The importance of our study is in reinforcing that both donor individuality and medium type have major impacts on PDX degradation, whilst the impact of temporal variation is limited. Of interest is that bifidobacterial growth was more stimulated in rich medium with PDX as carbon source, whereas growth of members of the *Bacteroidetes* were more stimulated in basal medium with PDX as carbon source. We recommend that variations in medium and donor samples should be considered when planning and interpreting *in vitro* incubation studies.

## INTRODUCTION

The human gastrointestinal (GI) tract is residence to trillions of microbes (mainly bacteria) that are commonly referred to as the “gut microbiota”, which is crucial to overall host health and disease (1, 2). The composition of the gut microbiota varies widely among individuals, and many factors such as host genetic background (3), antibiotic exposure (4), dietary habits (5), geography (6), and age (7) contribute to its diversity. Over the past decades, much progress has been achieved thanks to the development of molecular, mostly DNA-based approaches, with a phylogenetic framework provided by 16S ribosomal RNA (rRNA) gene sequencing (8), and functional insights unraveled by metagenomics (9, 10) as well as cultivation of pure and defined mixed (fecal) cultures (11). However, most *in vivo* studies are, in fact, a snapshot of which microbial groups are there, and how abundant they are. The lack of mechanistic insights often limits our understanding of the functional role of the different microbes. To this end, *in vitro* models can to some extent fill the gap between *in vivo* studies and single and defined mixed culture studies (11). Furthermore, *in vitro* models also allow the investigation of mechanistic effects of different dietary components on microbiota composition and metabolic activity independent of the host, making them cost-effective alternative approaches for gut microbiota research.

A range of *in vitro* models has been developed to mimic colonic fermentation, ranging from simple batch culture models to sophisticated multi-stage continuous culture models, and to artificial digestive systems (12). Among those models, the fecal batch culture (FBC) model is particularly employed to study production of short-chain fatty acids (SCFAs) and other metabolites from dietary compound fermentation by gut microbiota due to its low cost and simple settings (13). A recent study from our group reported good reflection of *in vivo* observations by FBC by comparing clinical intervention data and FBC data with respect to carbohydrate degradation (14). However, there are many choices to be made in the experimental set-up of FBCs that can influence compositional dynamics and activity of the microbiota that will be observed. For example, for running an FBC model study usually a single time-point feces sample is used, bringing challenges to the interpretation of results and the comparison between studies as variation within an individual over time could be sometimes substantial (15). Moreover, the well-established inter-individual variability in microbiota composition is often not taken into account as it is common practice to pool feces from different donors as inoculum (16, 17). In addition, the consequence of the choice of cultivation medium must not be underestimated as differences in e.g., nutrient availability can have a drastic effect on the selectivity of a given medium for microbial growth. This selectivity is not only determined by variations in nutrient composition, but also in nutrient load (18, 19). Last but not least, the methods used for sampling and cryopreservation of feces will determine which microbes will survive these procedures, notably when subjects are requested to collect samples at home. The gut microbiota contains a wide variety of microbes with different survival rates when exposed to oxygen varying from strict anaerobes to facultative anaerobes. Hence, several protocols have been developed to optimize these procedures in order to maximize recovery of viable microbes (20). Although complete preservation of all viable microbes in the original sample will be challenging, a recent study showed that fast introduction of anoxic conditions to fecal samples after defecation resulted in limited loss of viability (14).

In this study, our aim was to elucidate the impact of donor individuality, temporal variation in microbiota composition within an individual and culture medium type on microbiota composition and metabolic activity in an FBC setting. We used the fiber polydextrose (PDX), as model carbon and energy source as previous studies have shown that it stimulates the growth of a consortium of different microbes (21, 22), the composition of which differed between subjects (23). Using fecal microbiota from three healthy adult donors, this study aimed to answer the following three research questions: 1) to what extent are microbial activity (SCFAs, gaseous metabolites, pH) and microbiota composition in FBCs with PDX as carbon source affected by donor individuality, temporal variation and culture medium type; 2) which groups of bacteria are stimulated in a medium- and PDX-dependent manner; 3) how does microbial viability vary in inocula prepared from feces collected at different dates.

## MATERIALS AND METHODS

### Fecal material collection and storage

Fecal material was obtained from three healthy donors at three different days (**Fig. 1**). This study was exempted from medical ethical approval by the medical ethical committee of Wageningen University due to the low burden and risk for participants. Subjects with an intake history of antibiotics six months and pre- or probiotics one month prior to the first sampling were excluded. Three healthy young female adults were recruited. No dietary restrictions or recommendations were given prior to sampling. The donors provided their fecal material at two successive days (day 1 and 2) and one month later (day 30) for this study (**Fig. 1A**). An informed written consent was obtained from each donor. Each donor defecated into a user-friendly and hygienic stool collection device, the Fecotainer (Excretas Medical BV, Enschede, the Netherlands). To create anoxic conditions during transportation, Anaerocult^®^ A mini (Merck KGaA, Darmstadt, Germany) was activated with 10 mL nuclease-free water and placed inside the device. Together with the device, two opened AnaeroGen (AnaeroGen^TM^ 3.5 L Sachet, Thermo Scientific, Waltham, Massachusetts, US) bags were put into the transportation box (AnaeroPack™ 7.0 L Rectangular Jar, Thermo Scientific). Afterwards, the fecal material was transported to the laboratory within 3 h (14). The device was opened inside an anaerobic chamber filled with 96% nitrogen and 4% hydrogen. A portion of around 5 g feces was instantly frozen at −20 ℃ for microbiota composition analysis. Another 10 g feces was mixed with an anoxic solution consisting of 4 mL dialysate (Tritium Microbiologie, Eindhoven, the Netherlands), 19 mL nuclease-free water and 7 g glycerol, resulting in a 25% (w/w) fecal inoculum as previously described (24). The inoculum was then transferred into a serum bottle, sealed with a butyl rubber stopper and aluminum cap inside the anaerobic chamber, and stored at −80 ℃ (**Fig. 1B**).

**FIG 1.**
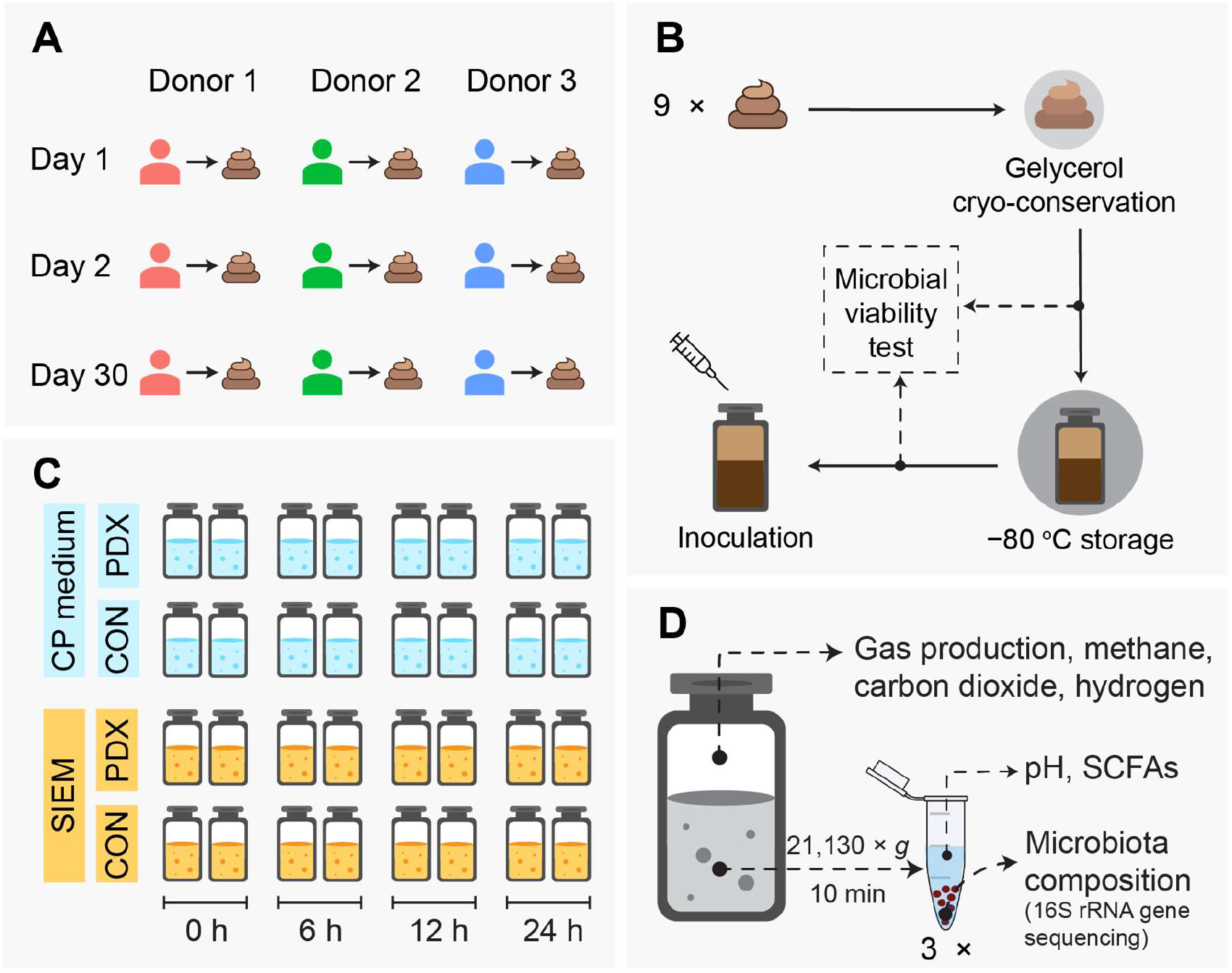
Schematic overview of the study set up. (A) feces collection, (B) glycerol cryopreservation and storage of fecal inoculum, concurrent with microbial viability test at pre- and post-storage, (C) *in vitro* incubation and (D) sample collection and measurement. CON: control, PDX: polydextrose, rRNA: ribosomal RNA, SCFAs: short-chain fatty acids, SIEM: standard ileal efflux medium.

### Microbial viability test

Before and after the storage at −80 ℃, 1 mL of each fecal inoculum was mixed with 2.5 µL of 20 mM propidium monoazide (PMA) dye (Biotium, Fremont, CA, USA). The samples were then incubated in the dark for 5 min, exposed to light in a PMA-Lite™ LED photolysis device (Biotium, Fremont, CA, USA) for light-induced crosslinking of PMA to dsDNA for 15 min at room temperature, and centrifuged at 4 ℃ at 5,000 × *g* for 10 min. After discarding the supernatant, the pellets were stored at −20 ℃ for DNA isolation and microbiota composition analysis.

### Culture medium preparation

Bicarbonate buffered CP medium (25, 26) served as the basal medium in this study. Per liter of medium, it consists of 0.41 g KH_2_PO_4_, 0.53 g Na_2_HPO_4_·2H_2_O, 0.3 g NH_4_Cl, 0.3 g NaCl, 0.1 g MgCl_2_·6H_2_O, 0.11 g CaCl_2_·2H_2_O, 4.0 g NaHCO_3_, 0.24 g Na_2_S·9 H_2_O, 0.5 g L-cysteine, 0.5 g resazurin, 1 mL acid stock solution (50 mM HCl, 1 mM H_3_BO_3_, 0.5 mM MnCl_2_, 7.5 mM FeCl_2_, 0.5 mM CoCl_2_, 0.1 mM NiCl_2_, 0.5 mM ZnCl_2_, 0.1 mM CuCl_2_), and 1 mL alkaline stock solution (10 mM NaOH, 0.1 mM Na_2_SeO_3_, 0.1 mM Na_2_WO_4_, 0.1 mM Na_2_MoO_4_). After boiling and cooling down, the headspace was exchanged to 80% nitrogen and 20% carbon dioxide. After autoclaving, 1 mL vitamin stock solution (20 mg/L biotin, 200 mg/L nicotinamide, 100 mg/L p-aminobenzoic acid, 200 mg/L thiamine (vitamin B1), 100 mg/L panthotenic acid, 500 mg/L pyridoxamine, 100 mg/L cyanocobalamine (vitamin B12), 100 mg/L riboflavin) was added through a 0.2 µm filter.

2-(N-morpholino) ethane sulfonic acid (MES) buffered standard ileal efflux medium (SIEM, Tritium Microbiologie B.V., Eindhoven, The Netherlands)) (27) served as the rich medium in this study. Per liter of medium, it consists of 50 ml BCO (60 g/L bacto peptone, 60 g/L casein, and 1 g/L ox bile), 16 mL salts solution (156.3 g/L K_2_HPO_4_, 281.3 g/L NaCl, 28.13 g/L CaCl_2_·2H_2_O, 0.31 g/L FeSO_4_ ·7H_2_O, 0.63 g/L hemin, 99% porcine), 10 mL 50 g/L MgSO_4_, 0.5 g L-cysteine, 1 mL vitamins (1 mg/L menadion, 2 mg/L D(+)-biotine, 0.5 mg/L Vitamin B12, 10 mg/L D(+) pantothenate, 5 mg/L aminobenzoic acid, 4 mg/L thiamine HCL and 5 mg/L nicotinamide adenine dinucleotide free acid), 0.5 g resazurin and 100 mL MES buffer (1M, pH = 6.0). Medium was prepared in the anaerobic chamber according to the manufacturer’s instructions.

### Carbon and energy source for incubations

PDX, kindly provided by Winclove Probiotics B.V. (Amsterdam, the Netherlands) was used as a carbon and energy source for incubations. It contains a minimal 90% of polymer and an average degree of polymerization of 12. PDX has a high solubility, allowing the preparation of 50% w/w solution at 20 ℃. Sterilized PDX was obtained by filtering the 50% w/w solution through a 0.2 µm filter.

### Inoculation and incubation

Inoculation and incubation were conducted in CP medium and SIEM, with PDX or nuclease-free water as control (CON) (**Fig. 1C**). Inoculation was carried out inside of the anaerobic chamber filled with 96% nitrogen and 4% hydrogen. 10 mL incubation vials were used in this study, whereby 5 mL medium was added to each vial. Cryo-conserved fecal inoculum was carefully thawed at 4 ℃ and added to the incubation vial at 0.1% (w/v) in duplicate. PDX was added at 1% (w/v). After completion, the vial was sealed with a butyl rubber stopper and aluminum cap inside the anaerobic chamber. For CP medium, the headspace gas was subsequently exchanged to 80% nitrogen and 20% carbon dioxide with a final pressure of 1.7 atmospheres. For SIEM, the headspace gas remained unchanged, i.e., 96% nitrogen and 4% hydrogen. Cultures were incubated in the dark at 37 ℃ statically for 24 h.

### Sample collection

Samples were collected at t = 0, 6, 12, and 24 h after inoculation (**Fig. 1C**). Specifically, at each time point, two incubation vials (biological replicate) in each group were used for sample collection (**Fig. 1D**). Firstly, the cumulative gas production along the incubation was determined by measuring the headspace gas pressure using GMH 3100 Series (GREISINGER, GHM Messtechnik GmbH, Regenstauf, Germany). Secondly, 0.2 mL headspace gas samples were taken for composition measurement. Thirdly, three aliquots of 1mL culture were then added into 1.5 mL sterilized Eppendorf tubes. After centrifuging at 4 ℃ at 21,130 × *g* for 10 min, the supernatant was separated from the pellet. One aliquot of the supernatant was used for pH measurement, the remaining two aliquots of supernatant and pellet were stored at −20 ℃ for SCFAs measurement and microbiota composition analysis, respectively.

### Metabolic activity parameters

Gas production was calculated with gas pressure (unit: psi) using the Ideal Gas Equation at 37 ℃ (28). Headspace gas composition including methane, carbon dioxide and hydrogen was measured using a CompactGC^4.0^ gas chromatograph (Global Analyser Solutions, Breda, The Netherlands), equipped with a thermal conductivity detector (TCD) and a Carboxen 1010 column. The settings for measuring were: pressure: 325 kPa, valve (injection) oven: 60 ℃, column oven temperature: 140 ℃, TCD temperature: 110 ℃, filament temperature 175 ℃, backflush time 30 s. Data were processed by the software Chromeleon (version 7.2.9, Thermo Fisher Scientific, Waltham, Massachusetts, USA). The concentrations of methane, carbon dioxide and hydrogen in the headspace were calculated based on the standard air pressure and the molar volume to convert the measured percentage to mmol/liter culture medium.

pH of the supernatant was measured by using ProLine Qis B210 pH meter (ProSense, Oosterhout, The Netherlands). Concentrations of SCFAs in the supernatant were measured via High-Performance Liquid Chromatography (HPLC, Shimadzu Prominence-I LC2030C-Plus, Shimadzu, Duisburg, Germany using a Shoedex SH1821 column (Showa Denko, New York, USA) and RID-20A refractive index detector (Shimadzu). Before injection, pre-treatment with Carrez A and B was carried out to de-proteinate the samples. Carrez A solution consists of 0.1 M K_4_Fe(CN)_6_·3H_2_O and Carrez B solution consists of 0.2 M ZnSO_4_·7H_2_O. Both solutions were stored at 4 ℃. Briefly, 500 µL of supernatant was mixed with 250 µL cold Carrez A, followed by mixing with 250 µL cold Carrez B. Next, the sample was centrifuged at 4 ℃ at 21,130 × *g* for 5 min, and the clear supernatant was collected for measurement. The conditions and settings of the HPLC instrument were as follows: eluent: 0.01 N H_2_SO_4_, eluent flow rate: 1 mL/min, column oven temperature: 54 ℃, flow rate: 0.8 mL/min, internal standard: 10 mM DMSO. All data were processed by software Chromeleon (version 7.2.9, Thermo Fisher Scientific).

### DNA isolation

Total DNA was isolated from fecal samples, PMA-treated inocula and collected FBC pellets following previously published protocol (29). Briefly, after resuspending samples with Stool Transfer and Recovery (STAR) buffer (Roche, Basel, Switzerland), the samples were repeatably bead beaten with beads in MP FastPrep-24 5^G^ (MP Biomedicals, Irvine, CA, USA) at 5.5 ms for 3 × 1 min. DNA was purified from the samples using the Maxwell 16 Tissue LEV Total RNA purification Kit custom-adapted for DNA extraction (cat no. AS 1220, Promega, Madison, Wisconsin, USA) in a Maxwell 16 MDx instrument (Promega). After purification, the DNA concentration was measured using NanoDrop 2000 (Thermo Scientific, Waltham, Massachusetts, USA).

### Quantitative PCR (qPCR) analysis

Quantification of the total bacterial 16S rRNA gene copy number was performed on the FBC pellet samples using the CFX384 Touch ™ Real-Time PCR Detection System (Bio-Rad, Hercules, California, USA). The primers used for amplification were 5’-TCCT ACGG GAGG CAGC AGT-3’ and 5’-GGAC TACC AGGG TATC TAAT CCTG TT-3’ (30). Each reaction mixture of 10 µL consisted of 5 µL BioRad iQ SYBR Green Supermix (BioRad), 0.2 µL forward primer (10 μM), 0.2 µL reverse primer (10 μM), 2.6 µL nuclease-free water and 2 µL template DNA (1 ng/µL). qPCR was performed using the following protocol: initial denaturation at 95 ℃ for 10 min, followed by 40 cycles of denaturation at 95 ℃ for 15 s, annealing at 60 ℃ for 30 s and elongation at 72 ℃ for 15 s. A melting curve from 60 ℃ to 95 ℃ in steps of 0.5 ℃ was added at the end. All qPCRs were performed in triplicates. Data were analyzed using Bio-Rad CFX Maestro software version 2.0.

### Microbiota composition analysis

Microbiota composition was determined by sequencing the V4 region of the 16S rRNA gene using Illumina Hiseq2500 technology, following the same procedure as described previously (14). Briefly, isolated and purified DNA was used for amplification of the V4 region of 16S rRNA gene with barcoded primers 515F (5’-GTGY CAGC MGCC GCGG TAA-3’) (31) and 806R (5’-GGAC TACN VGGG TWTC TAAT-3’) (32) in duplicate PCRs. PCR was performed in 50 µL system including 10 µL 5× HF buffer (Thermo Fisher Scientific, Vilnius, Lithuania), 1 µL dNTPs (10 mM, Thermo Fisher Scientific), 0.5 µL Phusion Hot start II DNA polymerase (2 U/µL, Thermo Fisher Scientific), 36.5 µL nuclease-free water (Promega, Madison, WI, USA), 1 µL DNA template (20 ng/µL) and 1 µL sample-specific barcoded primer (10 µM) (33). The PCR product was purified with the CleanPCR kit (CleanNA, The Netherlands), and quantified using the Qubit dsDNA BR Assay kit (Invitrogen by Thermo Fisher Scientific, Eugene, OR, USA). An equimolar mix of purified PCR products was prepared and sent to Novogene (Cambridge, United Kingdom) for sequencing. Raw sequence data were processed NG-Tax 2.0 with default settings (33, 34). Amplicon sequencing variant (ASV) picking and taxonomic assignments were performed using the SILVA 132 database.

### Statistical analysis

All statistical analyses were conducted in R (version 4.1.1). All figures were generated using *ggplot2* R package (35). 16S rRNA gene sequencing read counts were transformed to microbial relative abundance, as implemented in the *microbiome* R package (36). The absolute abundance of each taxon was calculated by multiplying the relative abundance with the total bacterial 16S rRNA gene copy number. Weighted UniFrac (37) and unweighted UniFrac (38) distance-based principal coordinates analysis (PCoA) was used to visualize microbial community variation at the ASV level. Permutational multivariate analysis of variance (PERMANOVA) was used to test for significant differences between groups, as implemented in the *vegan* R package (39). Furthermore, at each incubation time point, contribution of each explanatory variable (i.e., donor individuality, temporal variation, culture medium type and PDX supplementation) to the weighted and unweighted UniFrac based partitioning distance was assessed using the *adonis2* function in the *vegan* package. Redundancy analysis (RDA) was performed on SCFA concentrations using the *rda* function in the *vegan* package using stepwise forward and backward model selection. In addition, at each incubation time point, a permutation test was performed to estimate the contribution of each explanatory variable (i.e., donor individuality, temporal variation, culture medium type and PDX supplementation) to the overall variation of SCFA concentrations, using the *permu.hp* function in the *rdacca.hp* R package (40).

Differences with respect to metabolic activity parameter and absolute abundance of microbial taxa between groups were analyzed by fitting a linear mixed-effects model (LMM) to each dataset with donor individuality, temporal variation, culture medium type, PDX supplementation, and incubation time as the fixed effects and each duplicate as the random effect using the *lmer* function from *lmerTest* package (41). False discovery rate (FDR) correction based on Benjamini–Hochberg procedure was used to correct for multiple testing. In all cases, (adjusted) *P* values ≤ 0.05 were considered statistically significant.

### Data availability

Raw sequence data are available at the European Nucleotide Archive under the accession number PRJEB56217.

## RESULTS

Incubations of fecal microbiota from three different healthy adult donors collected at three different dates were carried out in two different media to elucidate the impact of donor individuality, temporal variation and culture medium type on microbiota composition and metabolic activity in FBCs with PDX as carbon and energy source. In addition, microbial viability in the inocula was determined to investigate its variation at different collection dates.

### Growth in two different media with or without PDX as carbon and energy source

CP medium served as the basal medium, while SIEM served as the rich medium in this study. As expected, in almost all CP medium incubations without PDX microbial growth was not observed as this medium does not contain any available energy source (**Fig. S1**). Only some growth was observed in the sample of donor 2 and this appeared to be limited to an unclassified member belonging to the family *Enterobacteriaceae*. Since CP medium without PDX does not provide an energy source for growth, we speculate this taxon grew on some residual energy source present in the respective fecal samples. Because of this very limited growth, samples in the control group (CON) with CP medium were excluded from all downstream analyses. Growth was observed in all other incubations, including SIEM without PDX. Growth in this medium was expected as SIEM contains bacto peptone and casein as nitrogen sources, which can also serve as energy sources for proteolytic microbes.

### Impact of donor individuality, temporal variation, culture medium type, and PDX supplementation on microbiota succession

PERMANOVA revealed a significant contribution of all examined variables to the microbiota succession in the FBCs. For unweighted UniFrac distances (presence and absence as well as phylogenetic relatedness of microbial taxa), the respective contributions were 34.4% for donor individuality, 21.1% for incubation time, 3.6% for PDX supplementation, 2.1% for culture medium type, and 2.0% for temporal variation, whereas these were 9.9%, 35.1%, 8.9%, 5.4% and 2.5%, respectively, based on weighted UniFrac distances (relative abundance and phylogenetic relatedness of microbial taxa). A clear separation between donor 1 and donors 2 & 3 was observed based on unweighted UniFrac distance calculations, while this was not the case for weighted UniFrac (**Fig. 2A, B**). The presence of methanogens (i.e., *Methanobrevibacter*) in donor 1 (**Fig. S2A**) explained the high contribution of donor individuality to the unweighted UniFrac distances as *Methanobrevibacter* is phylogenetically distant from all bacteria, and therefore its presence in some of the samples has a strong impact on the phylogenetic distance-based algorithm used in the unweighted UniFrac calculations.

**FIG 2.**
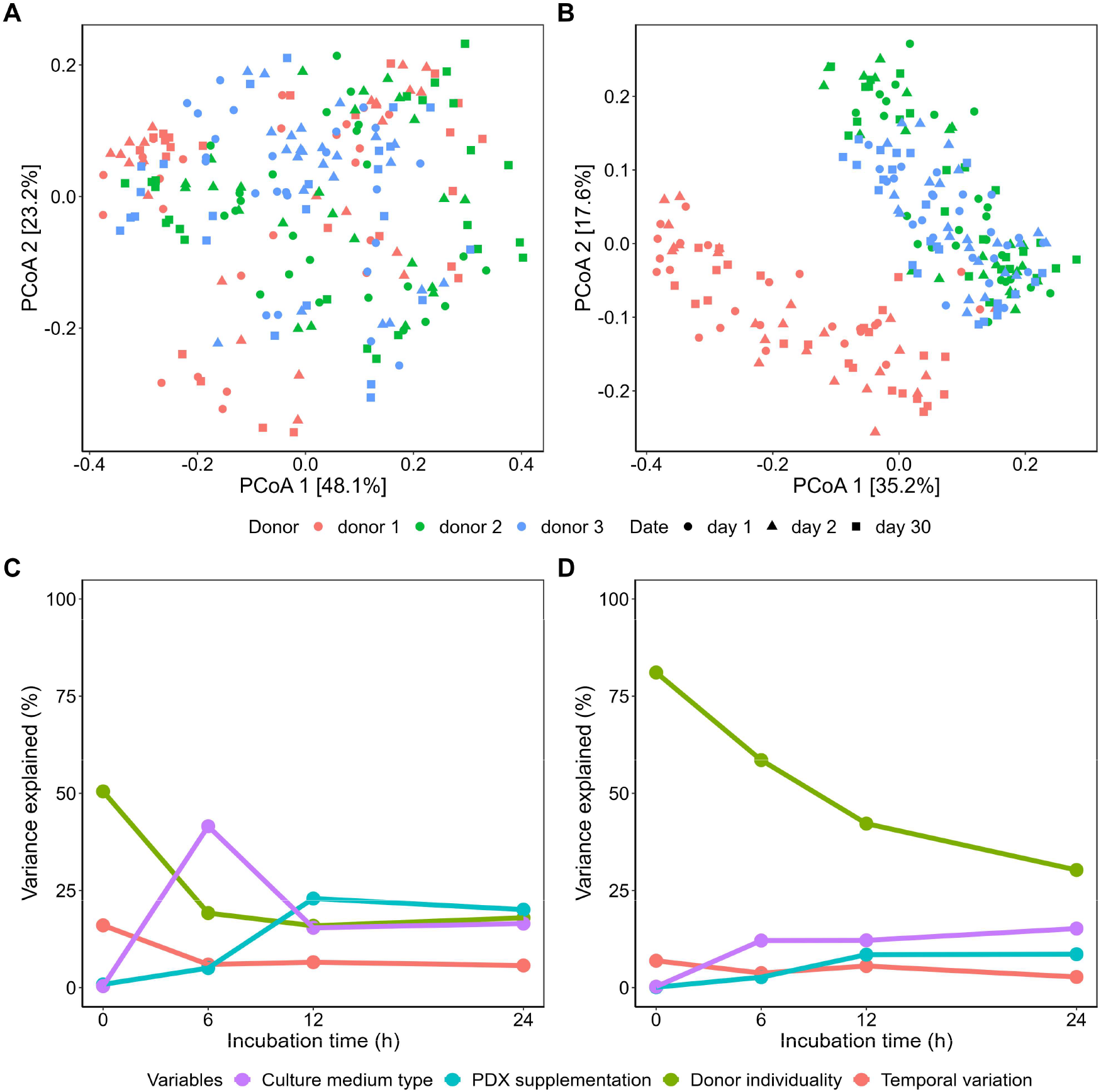
Microbiota composition variation during the incubation. PCoA of all samples based on (A) weighted UniFrac and (B) unweighted UniFrac distance matrices at ASV level. Variation of microbiota composition based on (C) weighted UniFrac and (D) unweighted UniFrac distance matrices that could be explained by donor individuality, temporal variation, culture medium type, or PDX supplementation at 0, 6, 12 and 24 h. ASV: amplicon sequencing variant, PCoA: principal coordinates analysis, PDX: polydextrose.

Specifically, at each incubation time point, donor individuality had a significant contribution (*P* = 0.001) to microbiota variation both for weighted UniFrac (50.49-15.90%) and unweighted UniFrac distances (81.10-30.27%), although the contribution decreased gradually during the incubation (**Fig. 2C, D**). With a similar trend but lower contribution, temporal variation could explain 16.02% and 6.91% of the variation based on weighted and unweighted UniFrac distances at 0 h, which decreased to 5.69% and 2.74%, respectively, at 24 h. In contrast, culture medium type had no contribution at 0 h as expected, but its contribution increased to 16.47% (weighted UniFrac) and 15.19% (unweighted UniFrac) at the end of the incubation (24 h). PDX supplementation also displayed increased contribution to 20.06% and 8.59% at 24 h based on weighted and unweighted UniFrac distances, respectively.

### Different microbial taxon groups observed during incubations

Due to donor individuality, and variations in media and carbon source, total microbial numbers as well as microbiota composition varied drastically between the incubations. Overall, the variation observed was reflected by 10 predominant genera across all incubations (**Fig. 3**).

**FIG 3.**
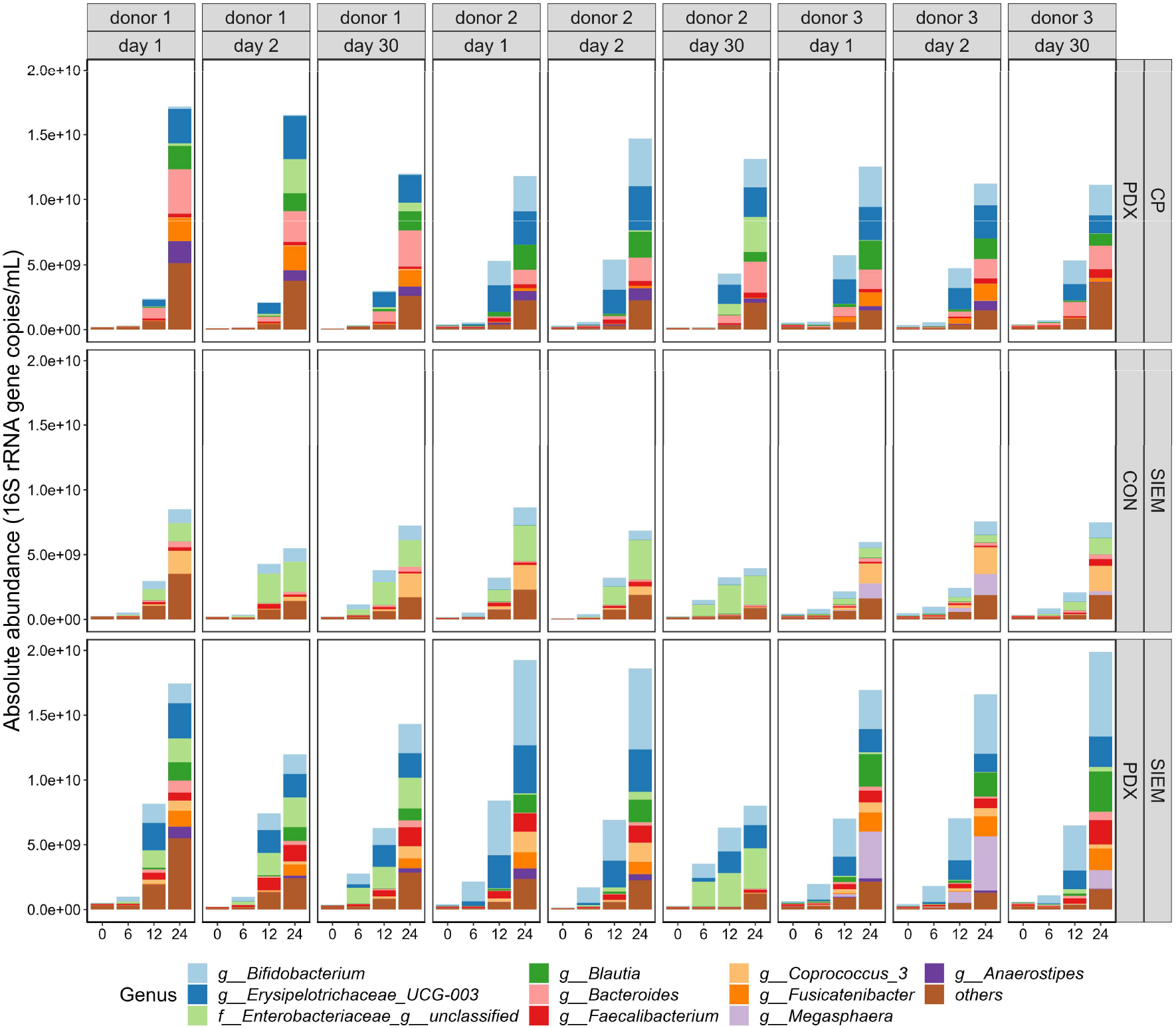
Microbial succession of the predominant microbial taxa during the incubation. Top 10 taxa at genus level were selected based on the ranking of the average absolute abundance across the entire dataset, other genera are summarised as “others”. Results from duplicates were averaged for visualization. CON: control, PDX: polydextrose, SIEM: standard ileal efflux medium.

When looking into the details of their growth individually, different patterns could be identified for specific groups of taxa by fitting a LMM to the their absolution abundance dynamics (**Fig. 4**, **Table S1**). Independent of whether growth occurred in CP medium or SIEM, in all subjects the genera *Erysipelotrichaceae* UCG-003, *Blautia*, and *Fusicatenibacter* were stimulated by PDX (**Fig.4**, **Table S1**, [LMM-PDX: coefficient > 0, *P* < 0.001], [LMM-SIEM: coefficient > 0, *P* > 0.05]), suggesting a medium-independent PDX stimulation manner. Although an increase in abundance of the genera *Anaerostipes* and *Bacteroides* was observed in all incubations, these taxa grew best in CP medium supplemented with PDX ([LMM-PDX: coefficient > 0, *P* = 0.008 and *P* = 0.613, respectively], [LMM-SIEM: coefficient < 0, *P* = 0.006 and *P* < 0.001, respectively]) whereas *Bifidobacterium*, *Faecalibacterium* and *Megasphaera* grew best in SIEM supplemented with PDX ([LMM-PDX: coefficient > 0, *P* < 0.001, *P* < 0.001 and *P* = 0.028, respectively], [LMM-SIEM: coefficient > 0, *P* < 0.001, *P* < 0.001 and *P* = 0.001, respectively]), suggesting a medium-dependent stimulation of PDX towards specific taxa. Remarkably, the genus *Megasphaera* was only observed in samples from donor 3, reinforcing the strong impact of individuality in microbial taxa at genus level on the incubations (**Fig. 4**).

**FIG 4.**
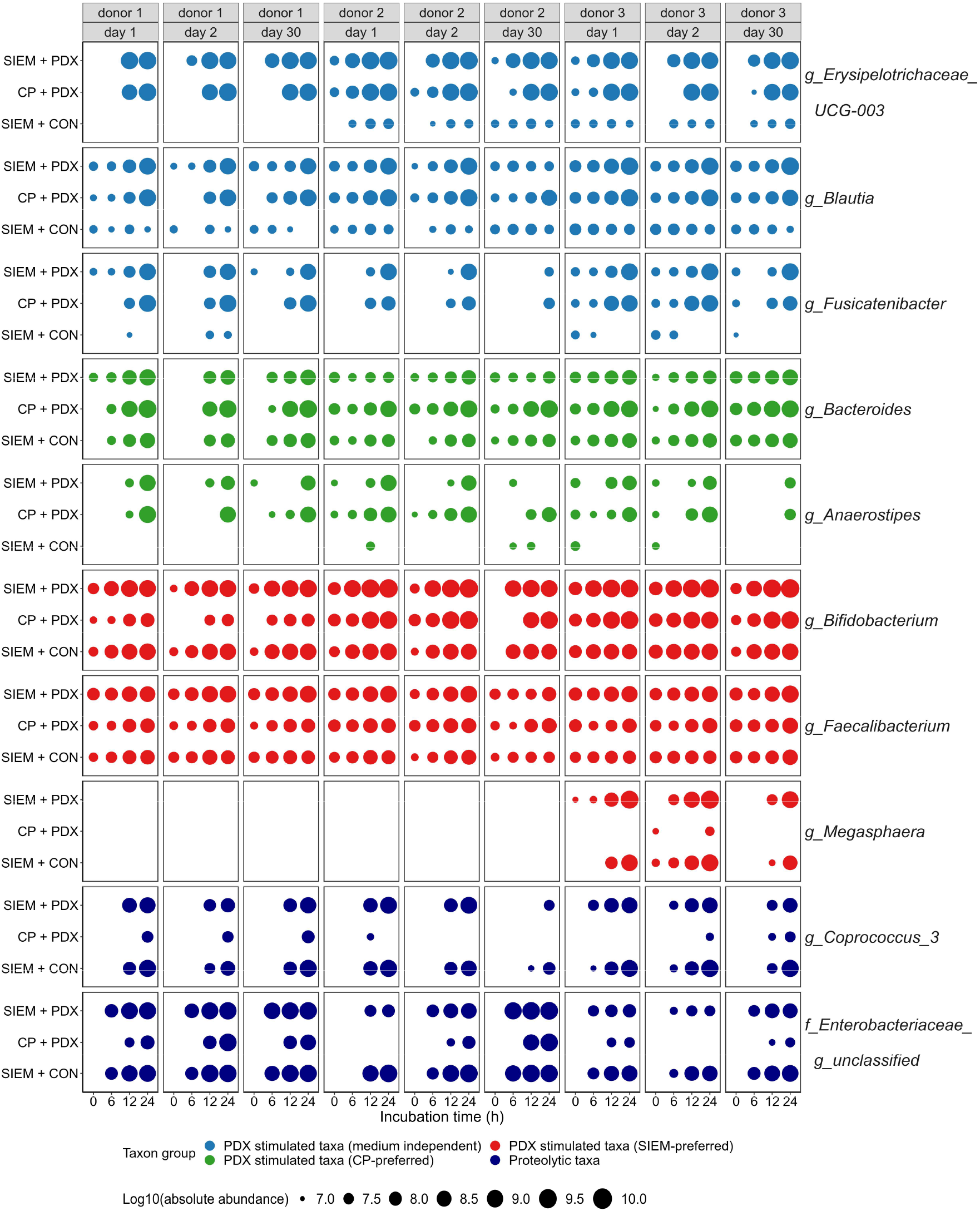
Dynamics of predominant microbial taxa during the incubation. Absolute abundances (log10 transformed) of each predominant taxon at genus level were plotted with the size of each circle representing the level, and the colour of each circle representing each taxon group. Blue: PDX stimulated taxa (medium-independent); green: PDX stimulated taxa (CP-preferred); red: PDX stimulated taxa (SIEM-preferred); navy-blue: proteolytic taxa. CON: control, PDX: polydextrose, SIEM: standard ileal efflux medium.

We did not observe any taxon that specifically grew in SIEM without PDX, but some taxa demonstrated better performance in SIEM compared to CP medium independent of the presence of PDX. One of the important differences between both media is the presence of ox-bile as well as of an organic nitrogen source, i.e., bacto-peptone and casein, suggesting that these taxa might engage in proteolytic activity. Taxa stimulated in SIEM included the genera *Coprococcus* and an unclassified genus within the family *Enterobacteriaceae* (**Fig.4**, **Table S1**, [LMM-PDX: coefficient < 0, *P =* 0.024 and *P* = 0.075, respectively], [LMM-SIEM: coefficient > 0, *P* < 0.001]).

### Impact of donor individuality, temporal variation, culture medium type, and PDX supplementation on metabolic activity variation

Incubation of fecal microbiota from three different donors collected at three different dates in two different media demonstrated different response to PDX with respect to the metabolic activity parameters. RDA revealed significant contribution of culture medium type (*P* = 0.012) and PDX supplementation (*P* = 0.002) to the overall variation in SCFA concentrations (**Fig. 5A**). RDA performed separately for different timepoints showed that the relative contribution of culture medium type gradually decreased after 6 h (from 51.59% at 6 h to 8.31% at 24 h), whereas contribution of PDX supplementation increased over time (from 9.33% at 6 h to 49.94% at 24 h) (**Fig. 5B**). Although the impact of these variables over time was similar to that observed for microbiota composition using the weighted UniFrac algorithm, PDX supplementation explained 50% of the variability on metabolic output at the end of the incubation, which contrasts to its relatively lower contribution to microbiota compositional observations (<20%) (**Fig. 2**).

**FIG 5.**
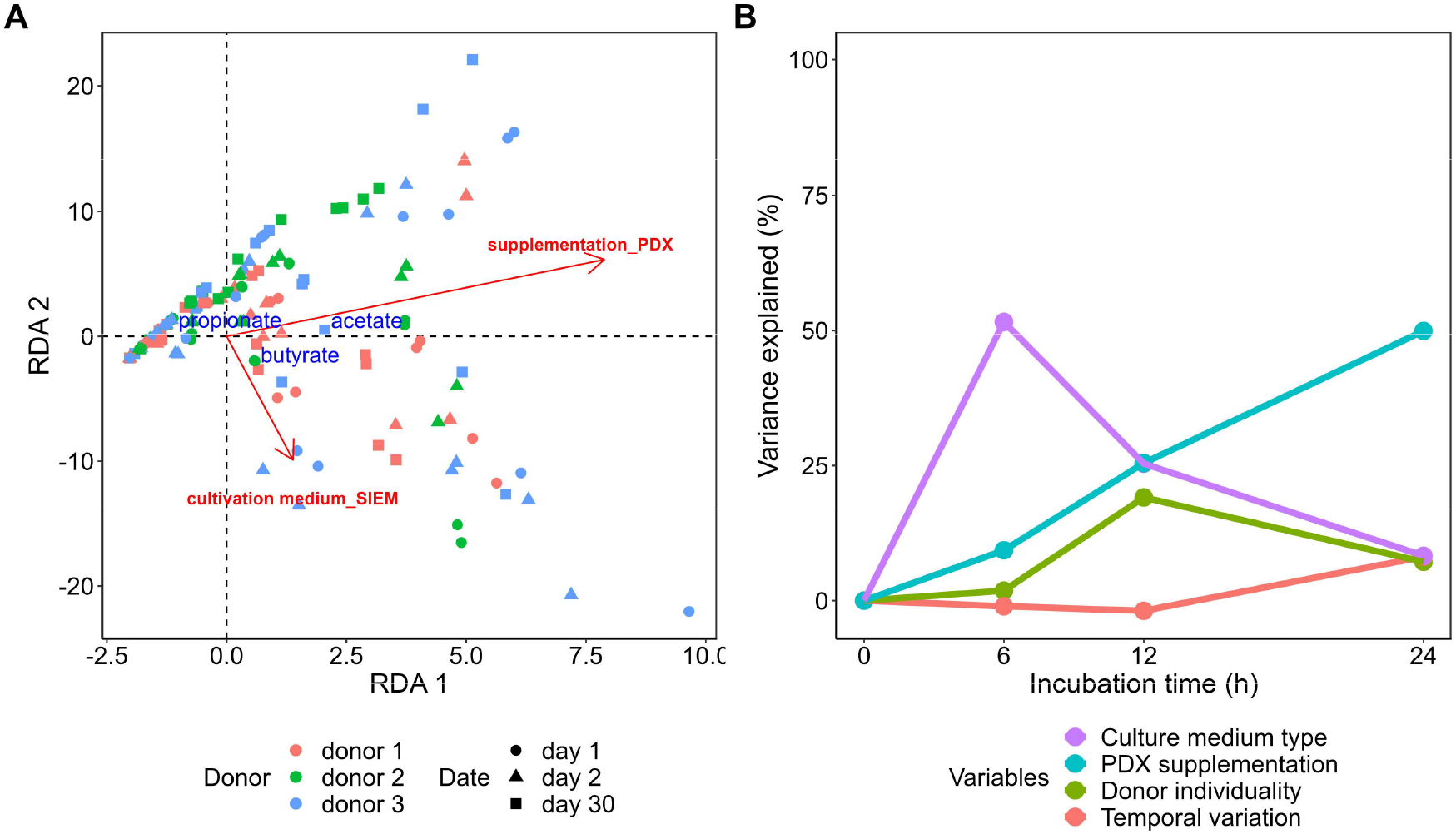
Variation of SCFA concentrations during the incubation. (A) RDA on the contribution of selected factors to the variation of SCFA concentrations. (B) Variation of SCFA concentrations that could be explained by donor individuality, temporal variation, culture medium type, or PDX supplementation at 0, 6, 12 and 24 h. Due to the difference between CP and SIEM medium with respect to pH and headspace composition, only SCFA data were included for the analysis. PDX: polydextrose, RDA: redundancy analysis, SCFAs: short-chain fatty acids, SIEM: standard ileal efflux medium.

By fitting a LMM to the metabolic activity parameter dynamics, the impact of those variables were tested (**Table S2**). Indeed, the supplementation of PDX promoted the production of all metabolites measured in this study, including acetate (+4.55 mmol/L, *P* < 0.001), propionate (+0.71 mmol/L, *P* < 0.001), butyrate (+2.05 mmol/L, *P* < 0.001), gas production (+64.30 ml/L, *P* = 0.004), carbon dioxide (+0.72 mmol/L, *P* = 0.008), hydrogen (+0.84 mmol/L, *P* < 0.001), and pH (−0.22, *P* < 0.001) (**Fig. S3**, **Fig. S4**, **Table S2**).

Culture medium type had a profound impact on all metabolic parameters measured in this study, except for propionate. Specifically, compared to CP medium, SIEM incubations showed significantly higher concentrations of acetate (+2.67 mmol/L, *P* < 0.001), butyrate (+1.67 mmol/L, *P* < 0.001), hydrogen (+1.55 mmol/L, *P* < 0.001), but significantly less gas (−146.90 ml/L, *P* < 0.001) (**Fig. S3**, **Fig. S4**, **Table S2**). Strikingly, hydrogen exclusively accumulated at 24 h in SIEM plus PDX incubations (**Fig. S4C**). Although carbon dioxide was also observed, a comparison between SIEM and CP could not be made as the latter one already contains carbon dioxide in the headspace as part of the carbonate buffer used with this medium. This difference in the buffer may also explain the significant lowering of pH (−3.02, *P* < 0.001) during SIEM incubations. No methane production was observed in any of the incubations despite the presence of the genus *Methanobrevibacter* in the samples of donor 1 (**Fig. S2A**).

A limited impact of donor individuality on metabolic parameters was observed over time, notably towards the end of the incubations (**Fig. 5B**). Compared to donor 1, it was found that donor 3 incubations had relatively higher acetate (**Fig. S3**, **Table S2**, +2.10 mmol/L, *P* < 0.001) and donor 2 had relatively lower propionate (−0.31 mmol/L, *P* = 0.034) concentrations. In line with the compositional observations, the impact of temporal variation was limited (<8.14%). Significantly lower concentrations of propionate (−0.48 mmol/L, *P* = 0.001) and butyrate (−1.23 mmol/L, *P* = 0.002) were found in samples from incubations with feces obtained on day 30 compared with day 1, mainly due to the lower microbial growth (**Fig. S1**) in incubations of fecal microbiota from donor 2 on day 30.

### Differences in viable microbiota composition of fecal inocula before and after cryopreservation

To determine the fraction of viable microbes in the inocula before and after −80 ℃ storage, microbial viability was determined with PMA, which selectively penetrates only into “dead” cells where it binds to double stranded DNA, inhibiting downstream PCR amplification. Because PMA does not enter viable cells, their DNA is amplified (42).

Before incubations were performed, fecal samples were collected, and subsequently, inocula were prepared and stored at −80 ℃ as glycerol stocks. To determine whether the storage at −80 ℃ impacted the viability of microbes and thereby may explain differences in observations between incubations, inoculum samples were treated with PMA before and after cryopreservation to determine the viable fraction. Weighted UniFrac distances revealed a significant difference with respect to relative abundance of viable microbiota after the cryopreservation (*P* = 0.039, **Fig. 6A**), although there was no significant impact on the presence and absence of viable microbes based on unweighted UniFrac distances (*P* = 0.288, **Fig. 6B**). At phylum level, the viable microbiota was predominated by *Firmicutes*, *Bacteroidetes* and *Actinobacteria* (**Fig. 6C**). The relative abundances of viable *Actinobacteria* (*P* = 0.039) and *Bacteroidetes* (*P* = 0.012) were increased whereas that of *Firmicutes* decreased (*P* = 0.008) (**Fig. 6D**). In addition, the fraction of viable *Actinobacteria* in the sample collected from donor 2 and donor 3 at day 30 was drastically lower compared to the other days, especially with respect to viable *Bifidobacterium* (**Fig. S2B**). Since *Methanobrevibacter* was observed in samples from donor 1, but methane production was not observed, we also checked its viability in the inocula (**Fig. S2A**). *Methanobrevibacter* was detected in all samples from donor 1 before and after PMA treatment, suggesting that loss of viability cannot explain the absence of methane. Although the absence of hydrogen in the CP medium incubations may explain the lack of methane production, we speculate for the SIEM incubations that there was no niche for methanogenic activity despite the production of hydrogen (**Fig. S4C**).

**FIG 6.**
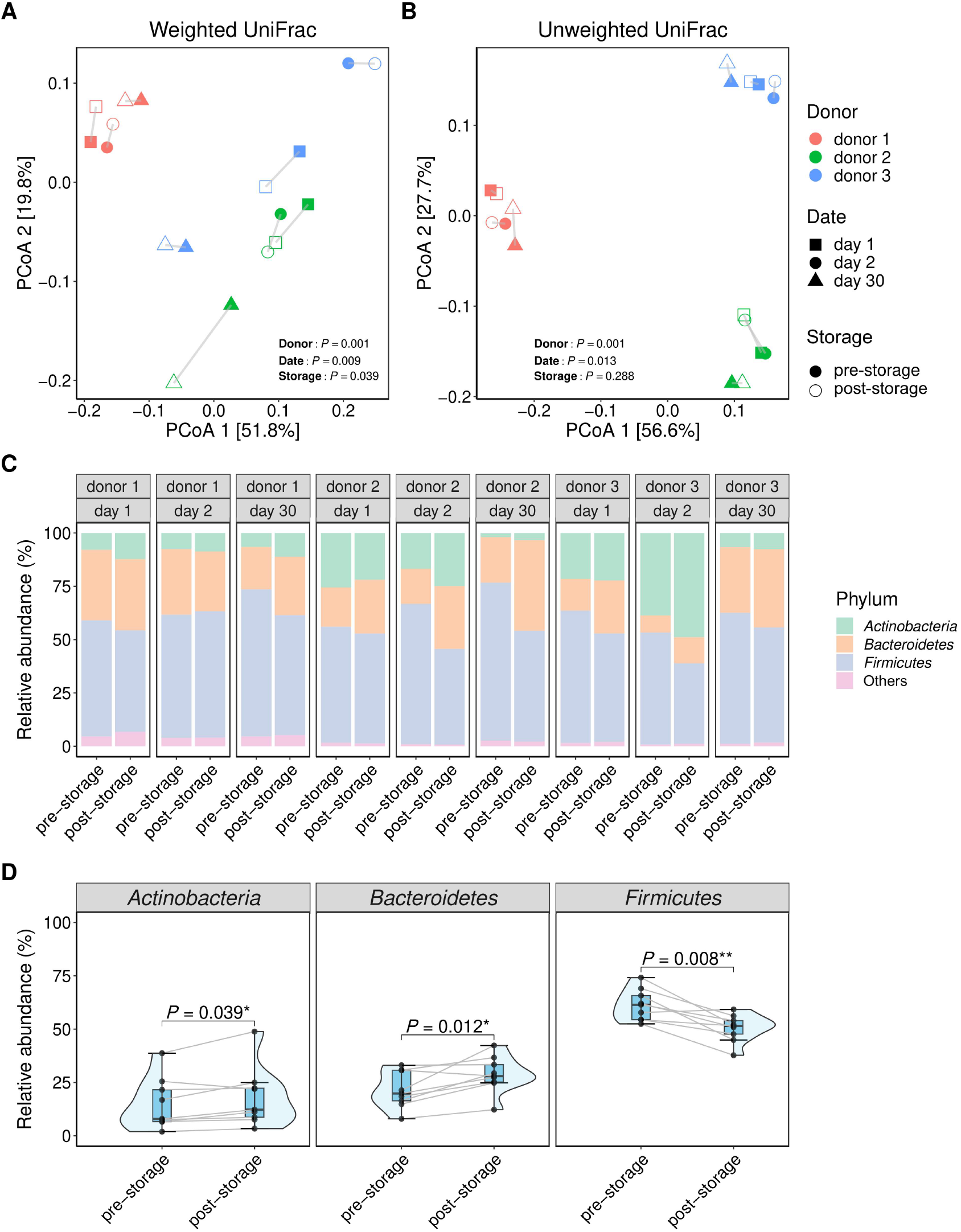
Impact of cryopreservation with glycerol at −80 ℃ on viable microbiota composition of fecal inocula. PCoA based on (A) weighted UniFrac and (B) unweighted UniFrac distances at ASV level. Pre- and post-storage samples are represented by solid and open symbols, respectively. Symbols are connected by lines per original sample. Significant differences in distances between groups of samples were assessed with PERMANOVA. (C) Relative abundance of taxa at phylum level and (D) changes of the predominant phyla in the fecal inocula after storage. Pre- and post-storage relative abundances of phyla were compared with paired Mann-Whitney *U* test. ASV: amplicon sequencing variant, PCoA: principal coordinates analysis, PERMANOVA: permutational multivariate analysis of variance.

## DISCUSSION

Using fecal microbiota from three donors, this study aimed to answer the following three research questions: 1) to what extent does donor individuality, temporal variation within the microbiota, and culture medium type explain the variation with respect to metabolite production and microbiota composition in FBCs with PDX as carbon source, 2) which groups of bacteria are stimulated in a medium- and PDX-dependent manner, 3) how does microbial viability vary in fecal inocula collected at different dates.

### Donor individuality rather than temporal variation determined microbiota composition in FBCs

With the restriction imposed by donor individuality, it has always been challenging to draw robust conclusions with respect to the fermentation of a variety of carbohydrates, even the succession of bacteria (43-45). Additionally, although gut microbiota has been considered as a stable ecosystem with strong resilience, the fluctuation caused by, e.g., diet changes, still exists, something that has been ignored in most *in vitro* studies (46). In our study, by partitioning the different factors that may contribute to microbiota succession, donor individuality was found to be a large determining factor of the variation of microbiota composition in FBCs. In agreement with our findings, An *et al.* (14) also found that subject individuality explained distinct portions of the overall microbiota composition variation, using the same FBC setting as used in this study, and investigating degradation of various carbohydrates with fecal microbiota from adults and elderly. For FBCs, the microbial composition of the fecal inoculum is the starting point for the microbial succession during incubation, and it also determines the overall microbiota composition over time. In our study, the presence of *Methanobrevibacter* was only observed in donor 1, and *Megasphaera* only in donor 3. This might partially contribute to the drastically distinct microbiota composition and dynamics. In another FBC study, it was also shown that the response of fecal microbiota to wheat bran and inulin was largely dependent on donor (47). These findings collectively indicated that donor individuality associated microbiota variation is an important contributing factor in FBC studies, which is lost when pooled fecal samples are used as inocula.

In contrast, temporal variation exerted limited impact, with less than 3% of variation attributed to it. Although the fluctuation of microbiota composition has been followed for example at the resolution of days (46) or weeks (48), relatively few studies took that into account while performing *in vitro* incubations. As found in our study, the fluctuation of microbiota over a short (i.e., one day) or longer period (i.e., one month) was minimal, and it had limited contribution to the variation of microbiota composition and SCFA production.

### PDX supplementation shaped microbial community and activity in a medium-dependent manner

SIEM has been widely used in studies with batch and continuous fermentation models of fecal microbiota as it represents well the proximal colon environment (e.g., lower pH, higher nutrition load compared to distal colon), where the fiber degradation starts. In comparison, CP medium is a basal medium developed mostly for the isolation and physiological characterization of pure cultures given its defined ingredient composition. In addition to that, higher pH and the lack of protein in CP medium also to some extent represent the conditions in the distal colon where most easily used substrates are scarce. As for the carbon and energy source, PDX was selected as it showed stimulation of different microbes in an individual-specific manner (21-23). In an *in vivo* situation, it was found that around 10% PDX (0.8 g out of 8 g consumption) ends up in the feces of healthy individuals (23). Additionally, a colon simulator study demonstrated a sustained degradation of PDX throughout the colon model (49). These findings indicate fermentation of PDX across the whole colon. Therefore, the combination of SIEM and CP medium provides complementary information, and thus can contribute to improved understanding of PDX fermentation by microbes, as carbohydrates degrading bacteria have species-specific preferences towards conditions such as pH and available energy source (50, 51).

Starting from 12 h, culture medium type and PDX supplementation also played important roles in the variation of microbiota composition. Despite the differences between media used in this study, most taxa stimulated by PDX were overlapping. Of note, a group of taxa (i.e., *Erysipelotrichaceae* UCG-003, *Blautia* and *Fusicatenibacter*) were dependent on PDX to grow, with limited growth in CON incubations with both media. However, culture medium type also exerted taxa-specific growth support with SIEM-preferred PDX stimulated taxa (e.g., *Bifidobacterium*) and CP medium-preferred PDX stimulated taxa (e.g., *Bacteroides*). The pH differences between CP medium and SIEM might contribute to the preference as growth of *Bacteroides* was favored in relatively higher (neutral) pH, while *Bifidobacterium* showed preference towards lower pH (51, 52). Additionally, the growth of *Bifidobacterium* spp. relies greatly on amino acids or peptides (53), which were available in SIEM rather than CP medium. Most *Bacteroides* spp. are capable of producing SCFAs via degrading complex carbohydrates (54). These findings collectively indicated that not only energy source determines which taxa successfully grow but also the medium/conditions in which they are offered.

Without PDX, nearly no bacterial growth or metabolic activity were observed in CP medium, whilst a group of genera (e.g., *Coprococcus*) could still grow in SIEM, regardless of the supplementation of PDX. This indicated that these genera are independent on PDX to grow. In agreement with our finding, *Coprococcus* was also previously found to be enriched in SIEM without any additional carbohydrates (14). In healthy individuals, the increase in *Coprococcus* was found to be positively associated with levels of branched-chain fatty acids (BCFAs, e.g., iso-butyrate and iso-valerate), which are the main end product of branched-chain amino acid (BCAA) degradation (55, 56). These findings collectively suggested *Coprococcus* is involved in protein degradation. Considering the nutrient differences between SIEM and CP medium such as peptone and casein in SIEM, this suggests that *Coprococcus* is involved in the proteolytic degradation of these substrates.

Although the microbiota composition as well as the compositional dynamics were found to be individual-specific in our study, metabolic activity, e.g., SCFA production showed no significant difference among incubations, indicating a donor-independent SCFA production manner. Similarly, by looking at the SCFA production from several substrates with fecal microbiota from three different donors at three different days, another FBC study showed that the amount and ratio of SCFAs were dependent on the type of substrate rather than the individual (57). In our study, SCFA production was explained best by culture medium type and PDX supplementation, with the highest contribution from the latter at 24 h. There is no doubt that the nutrients provided by the culture medium selectively dictate the metabolic activity in FBCs (18). Higher concentrations of SCFAs were found in the rich medium (i.e., SIEM) compared to the basal medium (i.e., CP medium) in our study. As the main end product of fiber degradation, SCFA concentrations were elevated with the supplementation of PDX as expected. Not only in *in vitro* models (58), colonic fermentation of PDX also induced higher SCFA production in a human intervention trial, notably acetate and butyrate (59).

In conclusion, our study demonstrated that donor individuality and medium choice impact the microbiota succession and metabolic activity during FBC when the same substrate is used as carbon and energy source. We recommend for future FBC studies to use individual fecal samples rather than pooling because within the same medium as we saw clear differences in microbiota composition when inoculated with the same substrate. In addition, we also recommend the use of basal and rich media to determine which microbes are stimulated by a certain substrate as these may reflect the variation in conditions along the colon as the proximal colon is richer in nutrients compared to the distal colon.

## ACKNOWLEDGEMENTS

We would like to thank all participants for providing the fecal donations. This work was funded by Winclove Probiotics B.V. (Amsterdam, The Netherlands). Zhuang Liu was furthermore financially supported by the China Scholarship Council (File No. 201806850091).

## AUTHOR CONTRIBUTIONS

The author contributions were as follows: conceptualization: Z.L., J.G., H.S., and E.G.Z.; funding acquisition: Z.L., J.G., H.S., and E.G.Z; investigation: Z.L.; writing original draft: Z.L..

## SUPPLEMENTARY FIGURES

**FIG S1.**
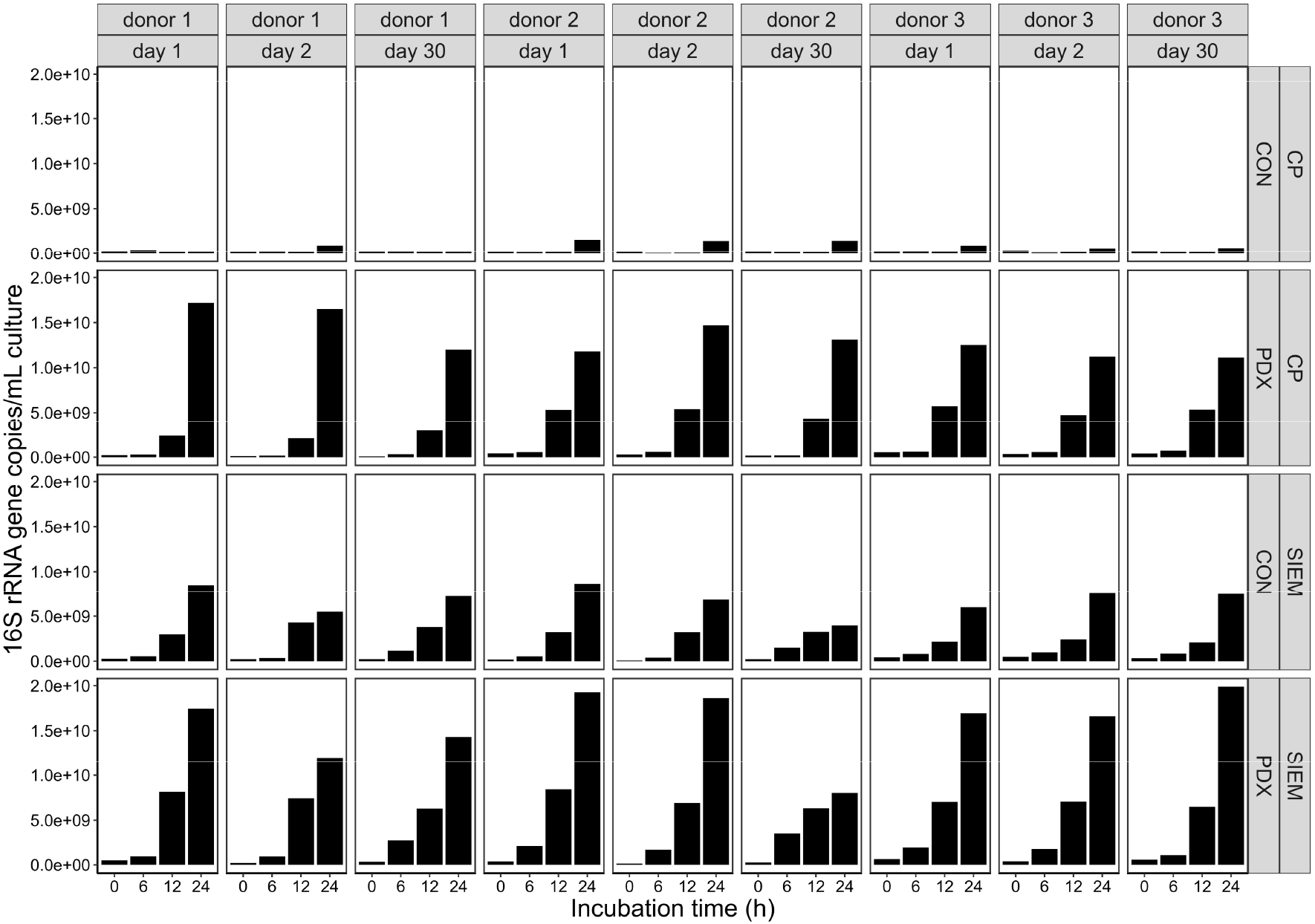
Total bacterial 16S rRNA gene copy numbers during the incubation. Results from the duplicates were averaged for visualization. CON: control, PDX: polydextrose, rRNA: ribosomal RNA, SIEM: standard ileal efflux medium.

**FIG S2.**
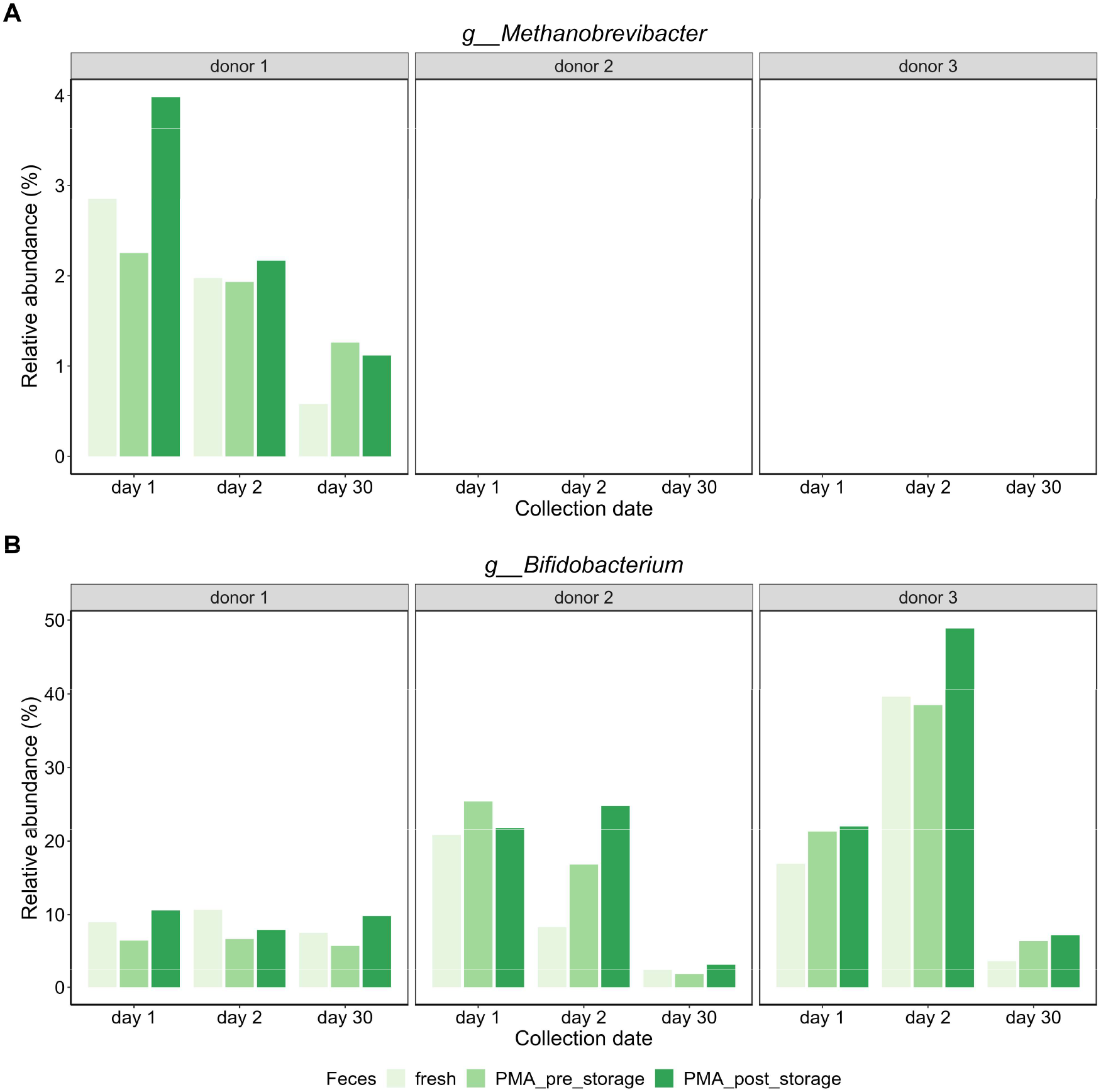
The relative abundance of (A) *Methanobrevibacter* and (B) *Bifidobacterium* in fresh feces, PMA-treated fecal inocula pre- and post-storage. PMA: propidium monoazide.

**FIG S3.**
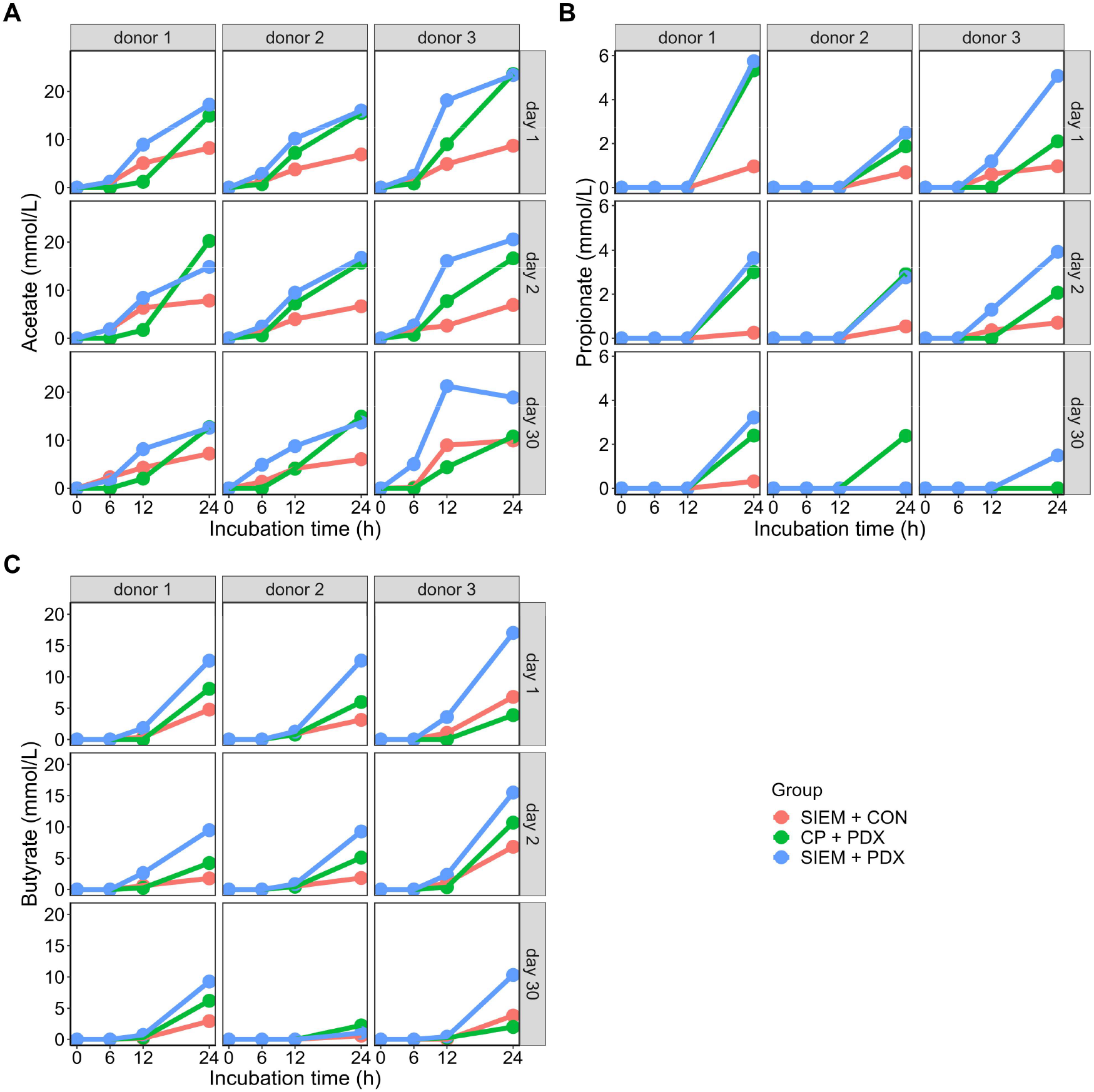
Concentration of (A) acetate, (B) propionate, and (C) butyrate in the cultures during the incubation. Results from the duplicates were averaged for visualization. Red lines represent (SIEM + CON), green lines represent (CP + PDX), and blue lines represent (SIEM + PDX). CON: control, PDX: polydextrose, SIEM: standard ileal efflux medium.

**FIG S4.**
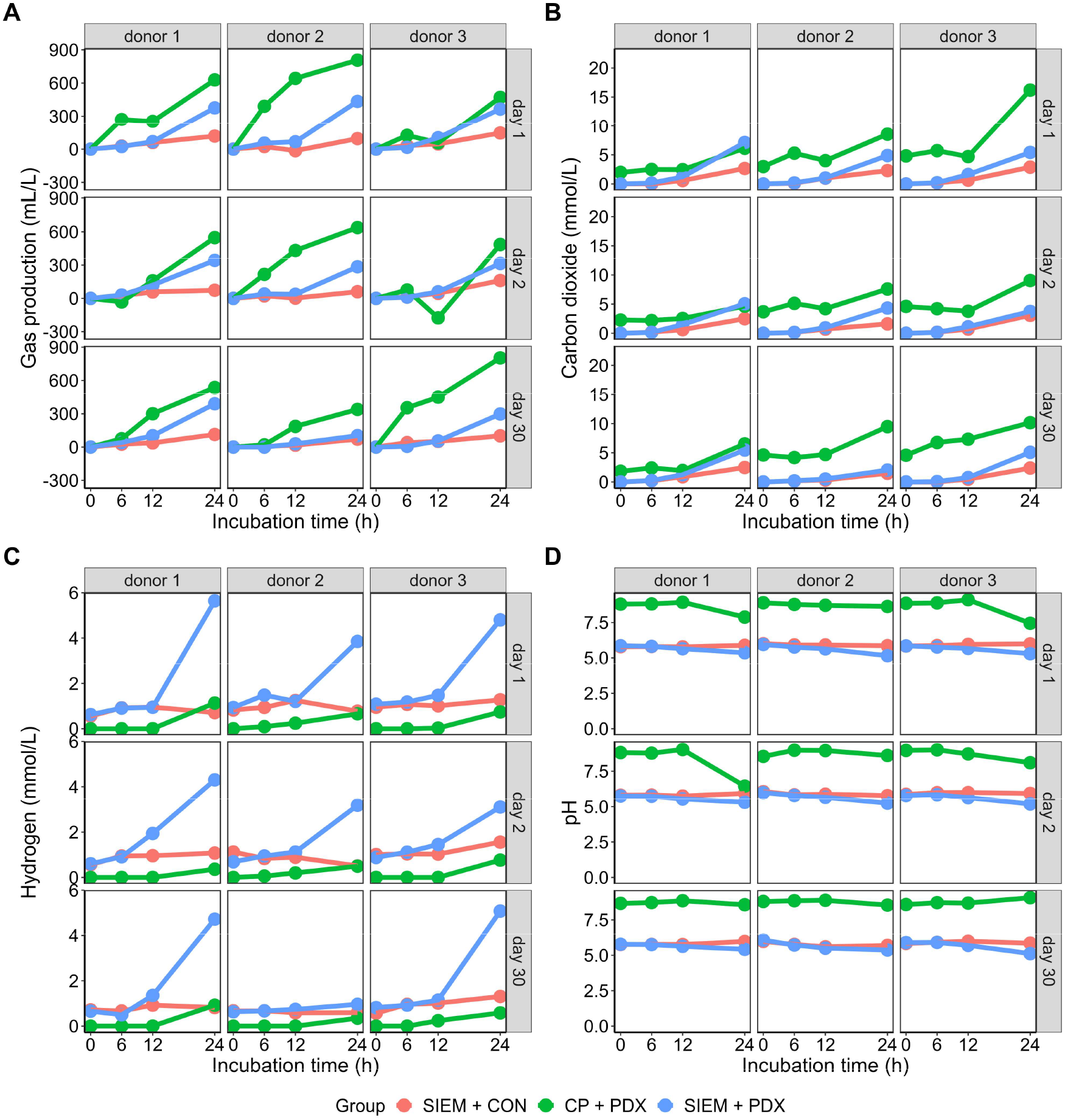
(A) Gas production, concentrations of (B) carbon dioxide, (C) hydrogen in the headspace, and (D) pH in the culture during the incubation. Results from the duplicates were averaged for visualization. Red lines represent (SIEM + CON), green lines represent (CP + PDX), and blue lines represent (SIEM + PDX). Methane was not observed in any of the incubations. CON: control, PDX: polydextrose, SIEM: standard ileal efflux medium.

## SUPPLEMENTARY TABLES

**TABLE S1.**
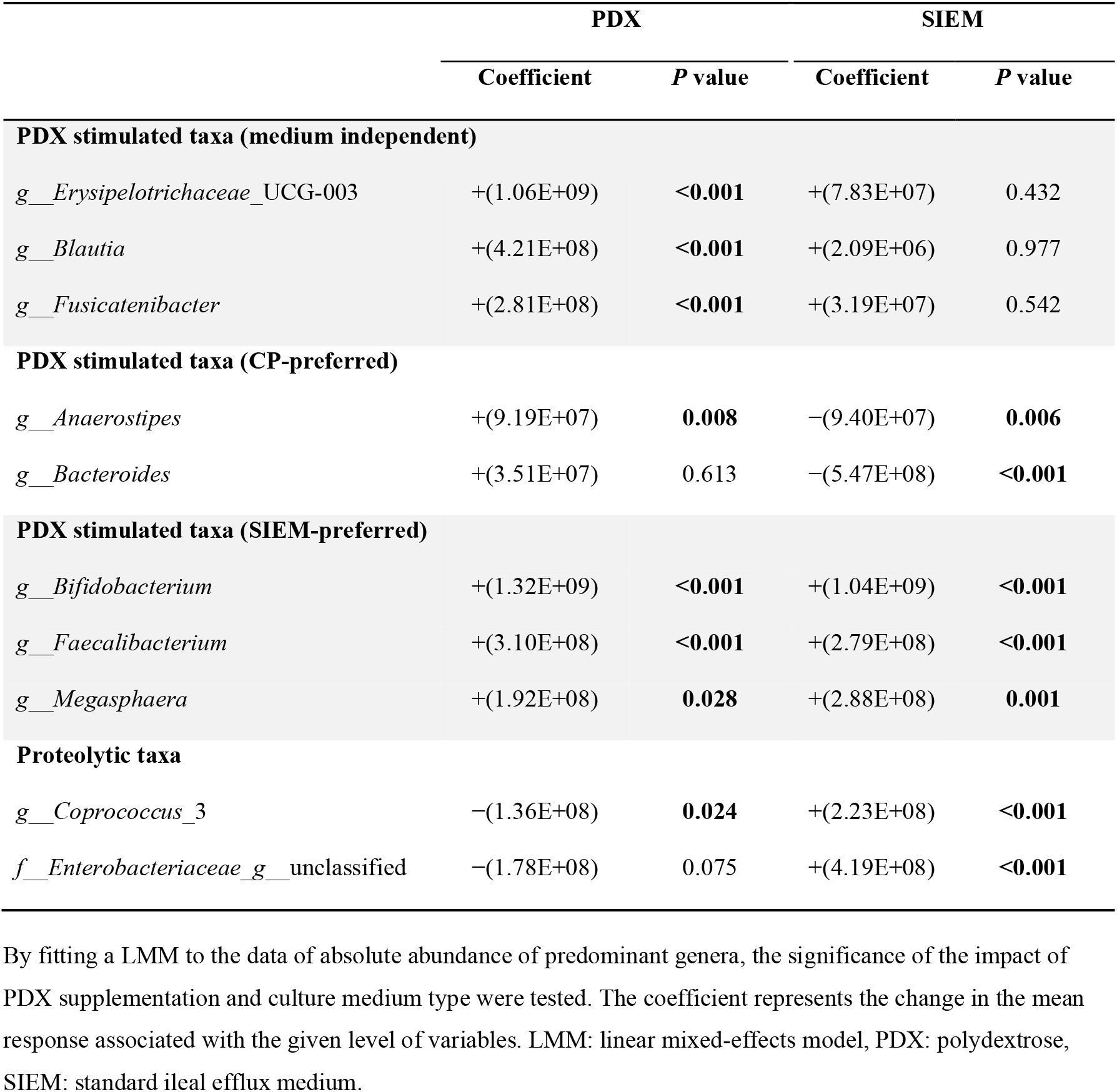
Significance test results by fitting a LMM to the data of absolute abundance of predominant genera.

|  | PDX |  | SIEM |  |
| --- | --- | --- | --- | --- |
|  | Coefficient | *P* value | Coefficient | *P* value |
| **PDX stimulated taxa (medium independent)** |  |  |  |  |
| *g__Erysipelotrichaceae_UCG-003* | +(1.06E+09) | **<0.001** | +(7.83E+07) | 0.432 |
| *g__Blautia* | +(4.21E+08) | **<0.001** | +(2.09E+06) | 0.977 |
| *g__Fusicatenibacter* | +(2.81E+08) | **<0.001** | +(3.19E+07) | 0.542 |
| **PDX stimulated taxa (CP-preferred)** |  |  |  |  |
| *g__Anaerostipes* | +(9.19E+07) | **0.008** | −(9.40E+07) | **0.006** |
| *g__Bacteroides* | +(3.51E+07) | 0.613 | −(5.47E+08) | **<0.001** |
| **PDX stimulated taxa (SIEM-preferred)** |  |  |  |  |
| *g__Bifidobacterium* | +(1.32E+09) | **<0.001** | +(1.04E+09) | **<0.001** |
| *g__Faecalibacterium* | +(3.10E+08) | **<0.001** | +(2.79E+08) | **<0.001** |
| *g__Megasphaera* | +(1.92E+08) | **0.028** | +(2.88E+08) | **0.001** |
| **Proteolytic taxa** |  |  |  |  |
| *g__Coprococcus_3* | −(1.36E+08) | **0.024** | +(2.23E+08) | **<0.001** |
| *f__Enterobacteriaceae_g__unclassified* | −(1.78E+08) | 0.075 | +(4.19E+08) | **<0.001** |

By fitting a LMM to the data of absolute abundance of predominant genera, the significance of the impact of PDX supplementation and culture medium type were tested. The coefficient represents the change in the mean response associated with the given level of variables. LMM: linear mixed-effects model, PDX: polydextrose, SIEM: standard ileal efflux medium.

**TABLE S2.** Significance test results by fitting a LMM to metabolic activity parameter data.

| Variables |  | Level | Coefficient | *P* value |
| --- | --- | --- | --- | --- |
| Acetate | Temporal variation | Day 2 | −0.32 | 0.537 |
|  |  | Day 30 | −0.73 | 0.161 |
|  | Donor individuality | Donor 2 | +0.40 | 0.440 |
|  |  | Donor 3 | +2.10 | **<0.001** |
|  | PDX supplementation | PDX | +4.55 | **<0.001** |
|  | Culture medium type | SIEM | +2.67 | **<0.001** |
| Propionate | Temporal variation | Day 2 | −0.15 | 0.293 |
|  |  | Day 30 | −0.48 | **0.001** |
|  | Donor individuality | Donor 2 | −0.31 | **0.034** |
|  |  | Donor 3 | −0.14 | 0.337 |
|  | PDX supplementation | PDX | +0.71 | **<0.001** |
|  | Culture medium type | SIEM | +0.24 | 0.096 |
| Butyrate | Temporal variation | Day 2 | −0.30 | 0.438 |
|  |  | Day 30 | −1.23 | **0.002** |
|  | Donor individuality | Donor 2 | −0.55 | 0.156 |
|  |  | Donor 3 | +0.55 | 0.156 |
|  | PDX supplementation | PDX | +2.05 | **<0.001** |
|  | Culture medium type | SIEM | +1.67 | **<0.001** |
| Gas production | Temporal variation | Day 2 | −45.51 | **0.041** |
|  |  | Day 30 | −31.37 | 0.158 |
|  | Donor individuality | Donor 2 | +6.54 | 0.768 |
|  |  | Donor 3 | −7.13 | 0.747 |
|  | PDX supplementation | PDX | +64.30 | **0.004** |
|  | Culture medium type | SIEM | −146.90 | **<0.001** |
| Carbon dioxide | Temporal variation | Day 2 | −0.47 | 0.086 |
|  |  | Day 30 | −0.25 | 0.357 |
|  | Donor individuality | Donor 2 | +0.47 | 0.085 |
|  |  | Donor 3 | +1.14 | **<0.001** |
|  | PDX supplementation | PDX | +0.72 | **0.008** |
|  | Culture medium type | SIEM | −3.59 | **<0.001** |
| Hydrogen | Temporal variation | Day 2 | −0.13 | 0.303 |
|  |  | Day 30 | −0.24 | 0.059 |
|  | Donor individuality | Donor 2 | −0.20 | 0.110 |
|  |  | Donor 3 | +0.08 | 0.524 |
|  | PDX supplementation | PDX | +0.84 | **<0.001** |
|  | Culture medium type | SIEM | +1.55 | **<0.001** |
| pH | Temporal variation | Day 2 | −0.03 | 0.515 |
|  |  | Day 30 | +0.02 | 0.701 |
|  | Donor individuality | Donor 2 | +0.10 | **0.039** |
|  |  | Donor 3 | +0.09 | 0.076 |
|  | PDX supplementation | PDX | −0.22 | **<0.001** |
|  | Culture medium type | SIEM | −3.02 | **<0.001** |

By fitting a LMM to the data of metabolic activity parameters, the significance of the impact of temporal variation, donor individuality, PDX supplementation and culture medium type were tested. The coefficient represents the change in the mean response associated with the given level of variables. LMM: linear mixed-effects model, PDX: polydextrose, SIEM: standard ileal efflux medium.

## REFERENCES

1. Backhed F, Ley RE, Sonnenburg JL, Peterson DA, Gordon JI. 2005. Host-bacterial mutualism in the human intestine. Science 307:1915–1920.

2. Gilbert JA, Blaser MJ, Caporaso JG, Jansson JK, Lynch SV, Knight R. 2018. Current understanding of the human microbiome. Nat Med 24:392–400.

3. Khachatryan ZA, Ktsoyan ZA, Manukyan GP, Kelly D, Ghazaryan KA, Aminov RI. 2008. Predominant role of host genetics in controlling the composition of gut microbiota. PLoS One 3:e3064.

4. Schwartz DJ, Langdon AE, Dantas G. 2020. Understanding the impact of antibiotic perturbation on the human microbiome. Genome Med 12:1–12.

5. David LA, Maurice CF, Carmody RN, Gootenberg DB, Button JE, Wolfe BE, Ling AV, Devlin AS, Varma Y, Fischbach MA, Biddinger SB, Dutton RJ, Turnbaugh PJ. 2014. Diet rapidly and reproducibly alters the human gut microbiome. Nature 505:559–563.

6. Yatsunenko T, Rey FE, Manary MJ, Trehan I, Dominguez-Bello MG, Contreras M, Magris M, Hidalgo G, Baldassano RN, Anokhin AP, Heath AC, Warner B, Reeder J, Kuczynski J, Caporaso JG, Lozupone CA, Lauber C, Clemente JC, Knights D, Knight R, Gordon JI. 2012. Human gut microbiome viewed across age and geography. Nature 486:222–227.

7. An R, Wilms E, Masclee AAM, Smidt H, Zoetendal EG, Jonkers D. 2018. Age-dependent changes in GI physiology and microbiota: time to reconsider? Gut 67:2213–2222.

8. Zoetendal EG, Cheng B, Koike S, Mackie RI. 2004. Molecular microbial ecology of the gastrointestinal tract: from phylogeny to function. Curr Issues Intest Microbiol 5:31–47.

9. Blottiere HM, de Vos WM, Ehrlich SD, Dore J. 2013. Human intestinal metagenomics: state of the art and future. Curr Opin Microbiol 16:232–239.

10. Cultrone A, Tap J, Lapaque N, Dore J, Blottiere HM. 2015. Metagenomics of the human intestinal tract: from who is there to what is done there. Curr Opin Food Sci 4:64–68.

11. Elzinga J, van der Oost J, de Vos WM, Smidt H. 2019. The use of defined microbial communities to model host-microbe interactions in the human gut. Microbiol Mol Biol R 83:e00054–18.

12. Payne AN, Zihler A, Chassard C, Lacroix C. 2012. Advances and perspectives in *in vitro* human gut fermentation modeling. Trends Biotechnol 30:17–25.

13. Macfarlane GT, Macfarlane S. 2007. Models for intestinal fermentation: association between food components, delivery systems, bioavailability and functional interactions in the gut. Curr Opin Biotech 18:156–162.

14. An R, Wilms E, Logtenberg MJ, Van Trijp MPH, Schols HA, Masclee AAM, Smidt H, Jonkers DMAE, Zoetendal EG. 2021. *In vitro* metabolic capacity of carbohydrate degradation by intestinal microbiota of adults and pre-frail elderly. ISME Commun 1:1–12.

15. Vandeputte D, De Commer L, Tito RY, Kathagen G, Sabino J, Vermeire S, Faust K, Raes J. 2021. Temporal variability in quantitative human gut microbiome profiles and implications for clinical research. Nat Commun 12:6740.

16. O’Donnell MM, Rea MC, O’Sullivan O, Flynn C, Jones B, McQuaid A, Shanahan F, Ross RP. 2016. Preparation of a standardised faecal slurry for ex-vivo microbiota studies which reduces inter-individual donor bias. J Microbiol Meth 129:109-116.

17. Aguirre M, Ramiro-Garcia J, Koenen ME, Venema K. 2014. To pool or not to pool? Impact of the use of individual and pooled fecal samples for in vitro fermentation studies. J Microbiol Meth 107:1–7.

18. Yousi F, Kainan C, Junnan Z, Chuanxing X, Lina F, Bangzhou Z, Jianlin R, Baishan F. 2019. Evaluation of the effects of four media on human intestinal microbiota culture in vitro. AMB Express 9:1-10.

19. Minnebo Y, Paepe D, Kim, Raes J, Wiele D, Van, Tom. 2021. Nutrient load acts as a driver of gut microbiota load, community composition and metabolic functionality in the simulator of the human intestinal microbial ecosystem. FEMS Microbiol Ecol 97:fiab111.

20. Hamilton MJ, Weingarden AR, Sadowsky MJ, Khoruts A. 2012. Standardized frozen preparation for transplantation of fecal microbiota for recurrent *Clostridium difficile* infection. Am J Gastroenterol 107:761–767.

21. Raza GS, Putaala H, Hibberd AA, Alhoniemi E, Tiihonen K, Mäkelä KA, Herzig K-H. 2017. Polydextrose changes the gut microbiome and attenuates fasting triglyceride and cholesterol levels in Western diet fed mice. Sci Rep 7:5294.

22. Roytio H, Ouwehand AC. 2014. The fermentation of polydextrose in the large intestine and its beneficial effects. Benef Microbes 5:305–313.

23. Costabile A, Fava F, Röytiö H, Forssten SD, Olli K, Klievink J, Rowland IR, Ouwehand AC, Rastall RA, Gibson GR, Walton GE. 2012. Impact of polydextrose on the faecal microbiota: a double-blind, crossover, placebo-controlled feeding study in healthy human subjects. Br J Nutr 108:471–481.

24. Maathuis A, Hoffman A, Evans A, Sanders L, Venema K. 2009. The effect of the undigested fraction of maize products on the activity and composition of the microbiota determined in a dynamic *in vitro* model of the human proximal large intestine. J Am Coll Nutr 28:657–666.

25. Plugge CM. 2005. Anoxic media design, preparation, and considerations. Environ Microbiol 397:3–16.

26. Stams AJ, Van Dijk JB, Dijkema C, Plugge CM. 1993. Growth of syntrophic propionate-oxidizing bacteria with fumarate in the absence of methanogenic bacteria. Appl Environ Microbiol 59:1114–1119.

27. Gibson GR, Cummings JH, Macfarlane GT. 1988. Use of a three-stage continuous culture system to study the effect of mucin on dissimilatory sulfate reduction and methanogenesis by mixed populations of human gut bacteria. Appl Environ Microbiol 54:2750–2755.

28. Theodorou MK, Lowman RS, Davies ZS, Cuddeford D, Owen E. 1998. Principles of techniques that rely on gas measurement in ruminant nutrition. BSAP Occas Publ 22:55–63.

29. Salonen A, Nikkila J, Jalanka-Tuovinen J, Immonen O, Rajilic-Stojanovic M, Kekkonen RA, Palva A, de Vos WM. 2010. Comparative analysis of fecal DNA extraction methods with phylogenetic microarray: effective recovery of bacterial and archaeal DNA using mechanical cell lysis. J Microbiol Meth 81:127–134.

30. Nadkarni MA, Martin FE, Jacques NA, Hunter N. 2002. Determination of bacterial load by real-time PCR using a broad-range (universal) probe and primers set. Microbiology 148:257–266.

31. Parada AE, Needham DM, Fuhrman JA. 2016. Every base matters: assessing small subunit rRNA primers for marine microbiomes with mock communities, time series and global field samples. Environ Microbiol 18:1403–1414.

32. Apprill A, Mcnally S, Parsons R, Weber L. 2015. Minor revision to V4 region SSU rRNA 806R gene primer greatly increases detection of SAR11 bacterioplankton. Aquat Microb Ecol 75:129–137.

33. Ramiro-Garcia J, Hermes GDA, Giatsis C, Sipkema D, Zoetendal EG, Schaap PJ, Smidt H. 2016. NG-Tax, a highly accurate and validated pipeline for analysis of 16S rRNA amplicons from complex biomes. F1000Res 5:1791.

34. Poncheewin W, Hermes GDA, Van Dam JCJ, Koehorst JJ, Smidt H, Schaap PJ. 2020. NG-Tax 2.0: a semantic framework for high-throughput amplicon analysis. Front Genet 10:1366.

35. Wickham H. 2016. ggplot2: elegant graphics for data analysis. Springer-Verlag New York.

36. Lahti L, Shetty S. 2017. microbiome R package.

37. Lozupone CA, Hamady M, Kelley ST, Knight R. 2007. Quantitative and qualitative β diversity measures lead to different insights into factors that structure microbial communities. Appl Environ Microbiol 73:1576–1585.

38. Lozupone C, Knight R. 2005. UniFrac: a new phylogenetic method for comparing microbial communities. Appl Environ Microbiol 71:8228–8235.

39. Oksanen J, Blanchet FG, Kindt R, Legendre P, Minchin P, O’Hara RB, Simpson G, Solymos P, Stevenes MHH, Wagner H. 2012. Vegan: community ecology package. R package version 2.0–2.

40. Lai J, Zou Y, Zhang J, Peres-Neto PR. 2022. Generalizing hierarchical and variation partitioning in multiple regression and canonical analyses using the rdacca.hp R package. Methods Ecol Evol 13:782–788.

41. Kuznetsova A, Brockhoff PB, Christensen RHB. 2017. lmerTest package: tests in linear mixed effects models. J Stat Softw 82:1–26.

42. Nocker A, Cheung CY, Camper AK. 2006. Comparison of propidium monoazide with ethidium monoazide for differentiation of live vs. dead bacteria by selective removal of DNA from dead cells. J Microbiol Meth 67:310–320.

43. De Paepe K, Verspreet J, Courtin CM, Van De Wiele T. 2020. Microbial succession during wheat bran fermentation and colonisation by human faecal microbiota as a result of niche diversification. ISME J 14:584–596.

44. Carlson J, Gould T, Slavin J. 2016. *In vitro* analysis of partially hydrolyzed guar gum fermentation on identified gut microbiota. Anaerobe 42:60–66.

45. Carlson J, Esparza J, Swan J, Taussig D, Combs J, Slavin J. 2016. *In vitro* analysis of partially hydrolyzed guar gum fermentation differences between six individuals. Food Funct 7:1833–1838.

46. Durbán A, Abellán JJ, Jiménez-Hernández N, Latorre A, Moya A. 2012. Daily follow-up of bacterial communities in the human gut reveals stable composition and host-specific patterns of interaction. FEMS Microbiol Ecol 81:427–437.

47. De Paepe K, Kerckhof FM, Verspreet J, Courtin CM, Van De Wiele T. 2017. Inter-individual differences determine the outcome of wheat bran colonization by the human gut microbiome. Environ Microbiol 19:3251–3267.

48. Faith JJ, Guruge JL, Charbonneau M, Subramanian S, Seedorf H, Goodman AL, Clemente JC, Knight R, Heath AC, Leibel RL, Rosenbaum M, Gordon JI. 2013. The long-term stability of the human gut microbiota. Science 341:1237439.

49. Mäkeläinen HS, Mäkivuokko HA, Salminen SJ, Rautonen NE, Ouwehand AC. 2007. The effects of polydextrose and xylitol on microbial community and activity in a 4-stage colon simulator. J Food Sci 72:M153–M159.

50. Belenguer A, Duncan SH, Holtrop G, Anderson SE, Lobley GE, Flint HJ. 2007. Impact of pH on lactate formation and utilization by human fecal microbial communities. Appl Environ Microbiol 73:6526–6533.

51. Walker AW, Duncan SH, Leitch ECM, Child MW, Flint HJ. 2005. pH and peptide supply can radically alter bacterial populations and short-chain fatty acid ratios within microbial communities from the human colon. Appl Environ Microbiol 71:3692–3700.

52. Duncan SH, Louis P, Thomson JM, Flint HJ. 2009. The role of pH in determining the species composition of the human colonic microbiota. Environ Microbiol 11:2112–2122.

53. Cui S, Gu Z, Wang W, Tang X, Zhang Q, Mao B, Zhang H, Zhao J. 2022. Characterization of peptides available to different bifidobacteria. LWT 169:113958.

54. Wexler HM. 2007. *Bacteroides*: the good, the bad, and the nitty-gritty. Clin Microbiol Rev 20:593–621.

55. Vijay A, Astbury S, Le Roy C, Spector TD, Valdes AM. 2021. The prebiotic effects of omega-3 fatty acid supplementation: A six-week randomised intervention trial. Gut Microbes 13:1863133.

56. Nestel N, Hvass JD, Bahl MI, Hansen LH, Krych L, Nielsen DS, Dragsted LO, Roager HM. 2020. The gut microbiome and abiotic factors as potential determinants of postprandial glucose responses: a single-arm meal study. Front Nutr 7:594850.

57. Mortensen PB, Hove H, Clausen MR, Holtug K. 1991. Fermentation to short-chain fatty acids and lactate in human faecal batch cultures. Intra- and inter-individual variations versus variations caused by changes in fermented saccharides. Scand J Gastroenterol 26:1285–1294.

58. Mäkivuokko H, Kettunen H, Saarinen M, Kamiwaki T, Yokoyama Y, Stowell J, Rautonen N. 2007. The effect of cocoa and polydextrose on bacterial fermentation in gastrointestinal tract simulations. Bioscience, Biotechnology, and Biochemistry 71:1834–1843.

59. Jie Z, Bang-Yao L, Ming-Jie X, Hai-Wei L, Zu-Kang Z, Ting-Song W, Craig SA. 2000. Studies on the effects of polydextrose intake on physiologic functions in Chinese people. Am J Clin Nutr 72:1503–1509.

